# Discriminating the Single-cell Gene Regulatory Networks of Human Pancreatic Islets: A Novel Deep Learning Application

**DOI:** 10.1101/2020.08.30.273839

**Authors:** Turki Turki, Y-h. Taguchi

## Abstract

Analyzing single-cell pancreatic data would play an important role in understanding various metabolic diseases and health conditions. Due to the sparsity and noise present in such single-cell gene expression data, analyzing various functions related to the inference of gene regulatory networks, derived from single-cell data, remains difficult, thereby posing a barrier to the deepening of understanding of cellular metabolism. Since recent studies have led to the reliable inference of single-cell gene regulatory networks (SCGRNs), the challenge of discriminating between SCGRNs has now arisen. By accurately discriminating between SCGRNs (e.g., distinguishing SCGRNs of healthy pancreas from those of T2D pancreas), biologists would be able to annotate, organize, visualize, and identify common patterns of SCGRNs for metabolic diseases. Such annotated SCGRNs could play an important role in speeding up the process of building large data repositories. In this study, we aimed to contribute to the development of a novel deep learning (DL) application. First, we generated a dataset consisting of 224 SCGRNs belonging to both T2D and healthy pancreas and made it freely available. Next, we chose seven DL architectures, including VGG16, VGG19, Xception, ResNet50, ResNet101, DenseNet121, and DenseNet169, trained each of them on the dataset, and checked prediction based on a test set. We evaluated the DL architectures on an HP workstation platform with a single NVIDIA GeForce RTX 2080Ti GPU. Experimental results on the whole dataset, using several performance measures, demonstrated the superiority of VGG19 DL model in the automatic classification of SCGRNs, derived from the single-cell pancreatic data.

## 1. Introduction

Recent advances in single-cell technologies have led to the generation of single-cell gene expression data, which help advanced computational methods to study the singlecell-derived gene regulatory networks (GRNs) [1-9]. By inferring GRNs, biologists would be able to get deeper insight into the working mechanism of cells, thereby improving their understanding of the mechanism underlying the regulation of cellular functions in various biological processes [10-12].

Zheng et al. [13] had proposed a computational learning approach, named scPADGRN, to infer dynamic gene regulatory networks (DGRN) consisting of several GRNs that change over time, using time-series single-cell data, as follows. First, *N* single-cell data (genes spanning over the rows and cells spanning over the columns) were provided, each associated with a different time point, followed by a cell-clustering step, to yield *N* cluster-specific data. The learning process aimed to infer DGRN by utilizing preconditioned ADMM for solving the optimization problem, where the objective function aimed to guarantee network precision, network sparsity, and network continuity of the DGRN. Next, subnetworks were extracted from DGRN, based on genes existing in the same specific pathways or biological processes. They proposed quantitative index, named DGIE, to predict regulatory relationships across the GRNs. Inferred GRNs were then compared against the ground truth of GRNs from Transcription Factor Regulatory Network database. Reported results for inferring DGRNs indicated scPADGRN to possibly help in the understanding of cell differentiation in different biological processes.

Prataba et al. [14] had evaluated the performance of 12 existing algorithms for inferring gene regulatory networks from different single-cell data, including simulated data from curated models and synthetic networks. Evaluation of algorithms was performed to assess the accuracy, stability, as well as network motifs and their stability. Aibar et al. had presented SCENIC to identify transcription factors regulating target genes for regulon identification and inference of gene regulatory networks. Van de Sande et al. [15] had presented a scalable version of SCENIC, to speed up the process of inferring GRNs.

Turki et al. [12] had proposed three machine learning-based approaches for the inference of singe-cell gene regulatory network (SCGRN) of T2D pancreas and healthy pancreas. The proposed ML approaches work under the supervised setting described as follows. The first ML approach (named FSL) provides a training set, consisting of positive and negative examples. Positive examples correspond to transcription factors (TFs) regulating target genes while negative examples correspond to the target genes that are not regulated by TFs. Feature vectors of the examples are constructed by the concatenation of expression of TFs and target genes. The training set is then provided to a machine learning algorithm, to induce a model. Feature vectors for test examples are constructed by considering concatenation of expression of TFs and target genes. Test examples are provided to the obtained model to perform prediction; the latter corresponds to +1 if a TF regulates a target gene, or -1 if a TF does not regulate a target gene. The second ML approach (named SSL) works similar to the FSL approach; the only difference is in the training and testing feature vector representations, in which a stacked autoencoder (SA) is applied to training and testing, to generate different feature vector representations. The resultant reduced training feature vectors from SA is concatenated with training feature vectors, and then provided to a machine learning algorithm to generate a model. The reduced testing feature vectors, obtained from applying SA to the test set, are concatenated with testing feature vectors. The trained model is then applied to the new testing feature vectors, to generate predictions. The third ML approach (named TSL) performs similar computations as in the SSL approach. However, instead of applying SA to the training and testing feature vectors, to get new feature representation concatenated with training feature vectors, the TSL applies SA to the extracted topological features, in order to be concatenated with training and testing feature vectors. Following similar steps as in the SSL approach, Iacono et al. [16] had proposed a computational approach for the inference of GRNs from single-cell T2D and healthy pancreas as well as from single-cell data of Alzheimer’s disease. Other methods have also been proposed to study the inference of GRNs [17, 18].

Compared to the recent approaches for inference of GRNs [12-16, 19], the current study is unique in several ways: (1) Previous methods had not taken full advantage of deep learning (DL) to improve the understanding of biological processes pertaining to cells. Since DL techniques have been successfully applied to solve many biological and medical problems, including image classification problems [20-29], we have presented a DL application for discriminating between GRNs, obtained from human pancreas islets, in a single-cell data. (2) We have generated an image dataset consisting of 224 GRNs from single-cell healthy and T2D pancreas; it has been made freely available in the supplementary *dataset*. (3) We have formulated the discrimination task as a binary classification problem to aid biologists in improving the visual experience while identifying healthy and disease-related GRNs. (4) We have utilized several DL architectures, such as VGG16 [30], VGG19 [30], Xception [31], ResNet50 [32], ResNet101[32], DenseNet121 [33], and DenseNet169 [33]. (5) We have performed extensive experimental evaluation to assess the feasibility of DL in tackling the proposed problem. (6) We have reported results that demonstrate the potential of DL in successfully tackling the problem of discriminating across SCGRNs. To the best of our knowledge, this work would set the foundation for advancing the biological understanding of various diseases and cellular mechanisms.

## 2. Material and Methods

### 2.1. Datasets of Healthy and T2D Pancreas

A chart showing the computational approach for generating an image dataset consisting of 224 single-cell-derived gene regulatory networks, pertaining to healthy and T2D pancreas, was prepared as follows (Figure 1). First, the human pancreas dataset was obtained from Segerstolpe et al. [34], and single-cell data were obtained from Iacono et al. [16, 19]. Single-cell gene regulatory networks were inferred for both healthy and T2D pancreas, as reported by Iacono et al. [16, 19]. Finally, we visualized gene regulatory networks using NetBioV package [35] in R, as reported previously [12]. Different layout styles were utilized in NetBioV, yielding 112 SCGRNs of healthy pancreas and 112 SCGRNs of T2D pancreas. Some SCGRN image samples are shown in Figure 2. The first row corresponds to 3 SCGRNs of healthy pancreas donors with distinct parameters while the second row corresponds to 3 SCGRNs of T2D pancreas donors with distinct parameters. All 224 image data generated are available in the supplementary *dataset*.

**Figure 1:**
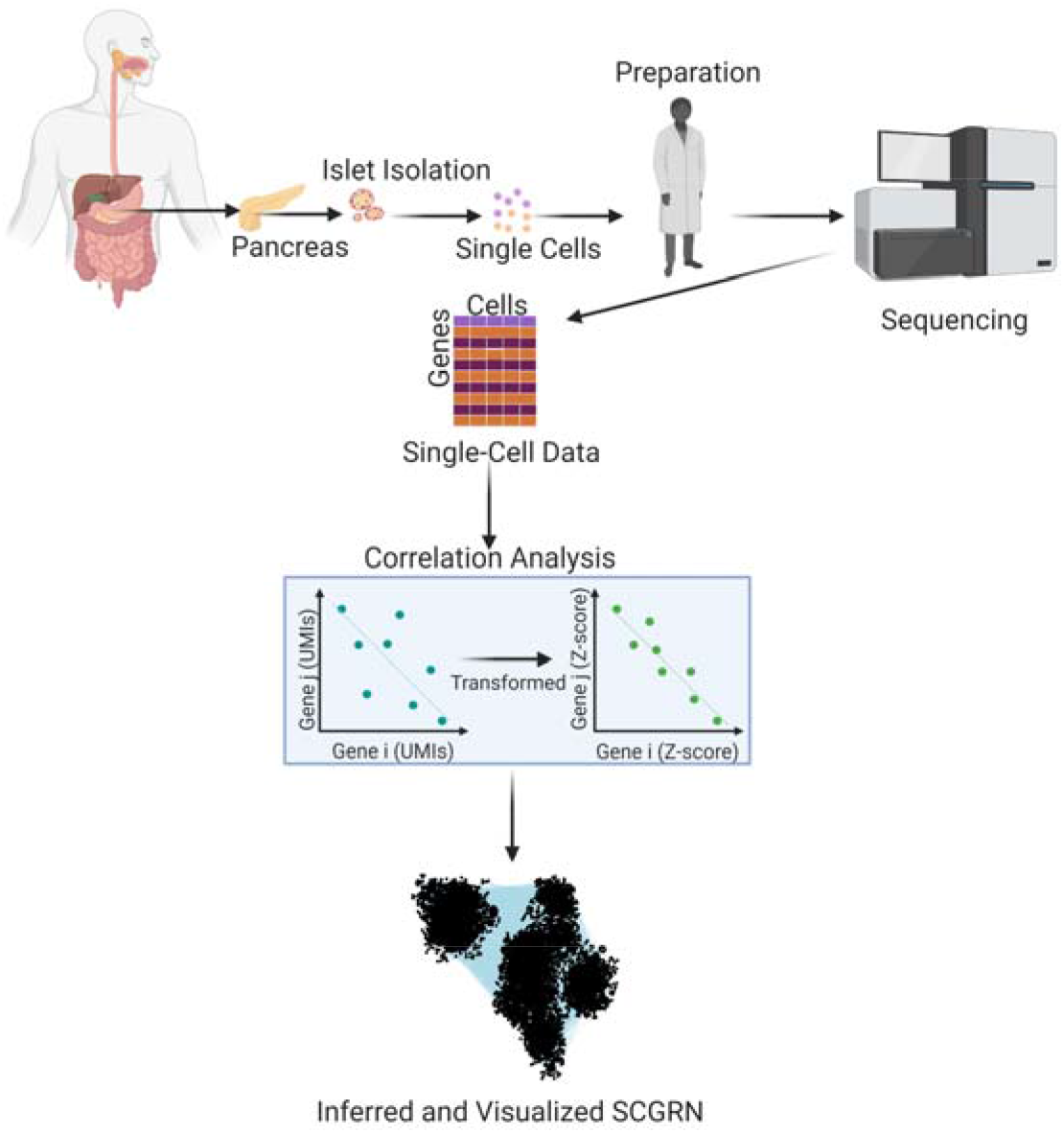
An overview of the computational approach for SCGRN image generation. Figure created with Biorender.com.

**Figure 2:**
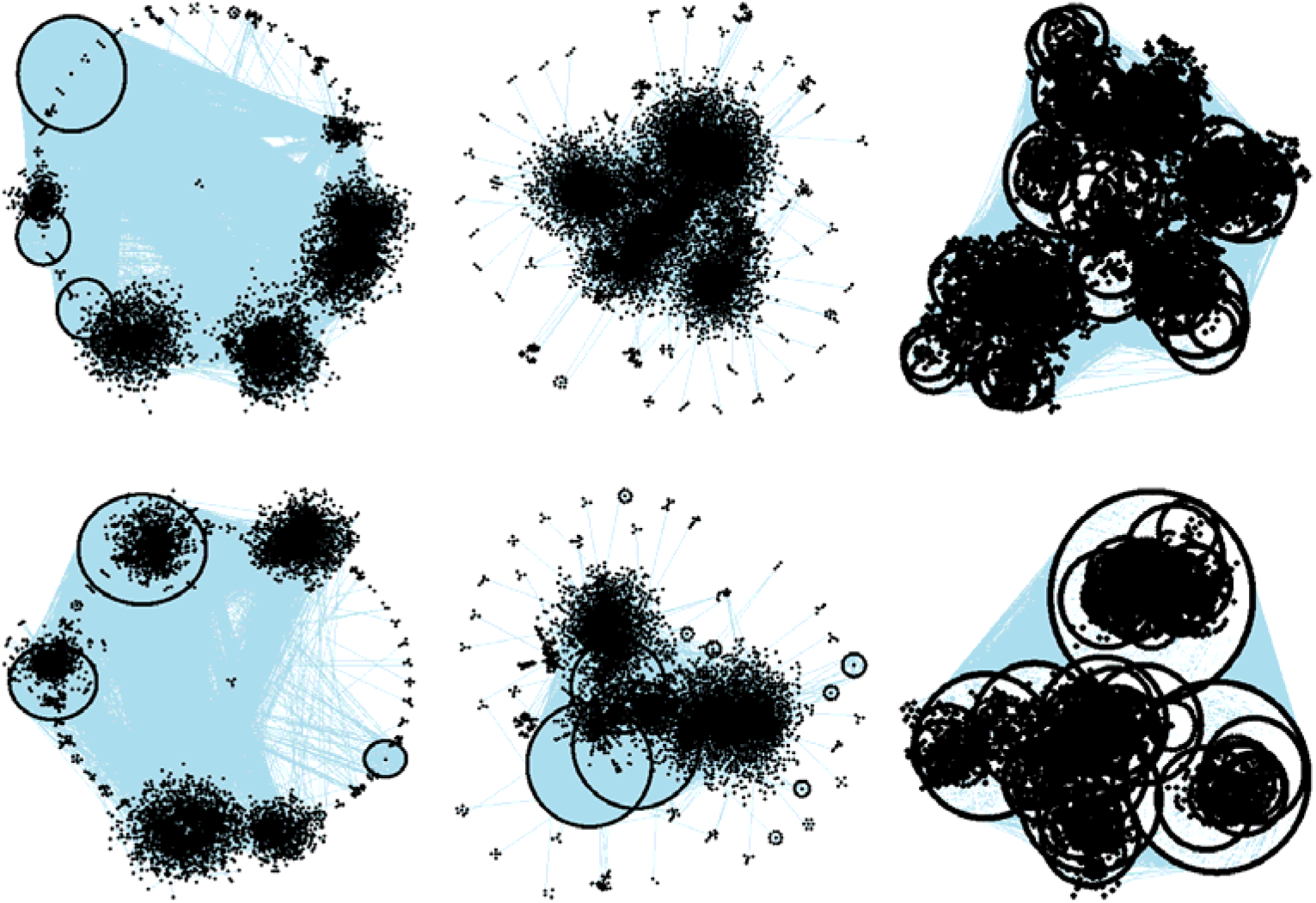
Samples of generated SCGRN image data. The top row corresponds to three SCGRNs of healthy pancreas while the bottom row corresponds to SCGRNs of T2D pancreas.

### 2.2. DL Framework

The computational framework of utilizing DL for discriminating between SCGRNs of healthy and T2D pancreas is shown in Figure 3. Suppose we are given a training set 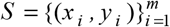 consisting of *m* labeled SCGRN images, along with a testing example of an unseen SCGRN image. The class label (1 or 0) of each training example is known, 1 representing an SCGRN pertaining to healthy pancreas and 0 representing an SCGRN corresponding to T2D pancreas. In this study, we utilized seven deep convolutional neural network architectures, including VGG16, VGG19, Xception, ResNet59, ResNet101, DenseNet121, and DenseNet169. The only difference was that we replaced the densely connected classifier in each pre-trained architecture by a densely connected classifier consisting of two dense layers. The first dense layer used ReLu activation, followed by a dropout. Since we aimed to perform binary classification, the second dense layer had 1 unit and a sigmoidal activation. Details for each architecture are provided in supplementary *models* file. For each architecture, we trained DL model on a training set of SCGRN images. Then, we applied the trained model to test examples consisting of images pertaining to SCGRNs of both healthy and T2D pancreas, in order to generate predictions corresponding to probabilities mapped to 1 (i.e., healthy pancreas), if the predicted probability is greater than 0.5; otherwise, it was mapped to 0 (i.e., T2D pancreas).

**Figure 3:**
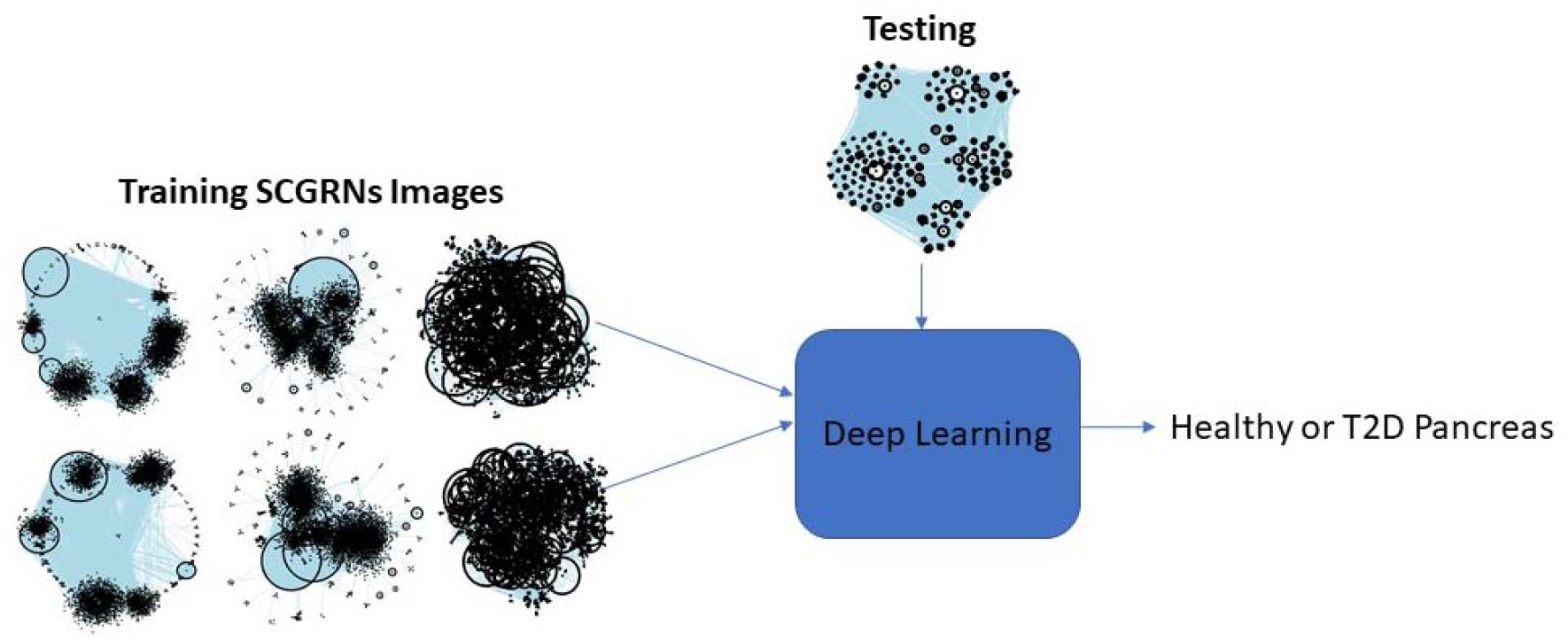
An illustration of the deep learning framework utilized in this study for distinguishing between SCGRNs of human pancreatic islets in healthy individuals and patients with T2D.

## 3. Experimental Study

### 3.1. Classification Methodology

We explored the behavior of seven DL architectures—VGG16, VGG19, Xception, ResNet50, ResNet101, DenseNet121, and DenseNet169—for inferring SCGRNs of single-cell pancreatic data of healthy and T2D pancreas. All architectures worked under supervised setting, where a training set was provided to train each DL model. The trained DL models were next applied to the test set to generate predictions mapped to 1 (healthy pancreas) or 0 (T2D pancreas). We utilized four different performance measures: Accuracy (ACC), F1, Matthews correlation coefficient (MCC), and area under curve (AUC). These measures were calculated based on the confusion matrix in Table 1 as follows:

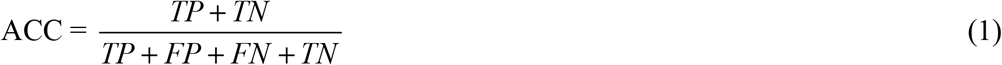

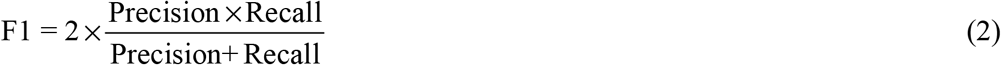

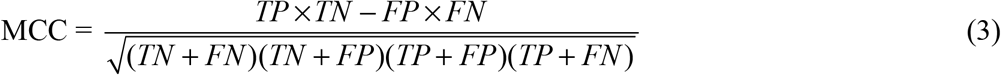

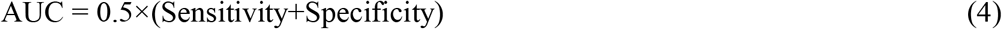

**Table 1:** Two-class confusion matrix.

| ACTUAL CLASS | PREDICTED CLASS |  |  |
| --- | --- | --- | --- |
|  |  | Class = H (1) | Class = T2D (0) |
|  | Class = H (1) | TP (True Positive) | FN (False Negative) |
|  | Class = T2D (0) | FP (False Positive) | TN (True Negative) |

For validation, we utilized 5-fold cross-validation mode as follows. We partitioned the SCGRN image datasets into 5 splits. In fold-1, we assigned 4 splits to training set, in order to train DL models. Then, we performed a prediction on 1 test split. Such a process was repeated for the other 4 folds. As can be seen from Figure 4, splits colored in black covered the whole dataset. Therefore, such validation showed the performance of DL models on the whole dataset.

**Figure 4:**
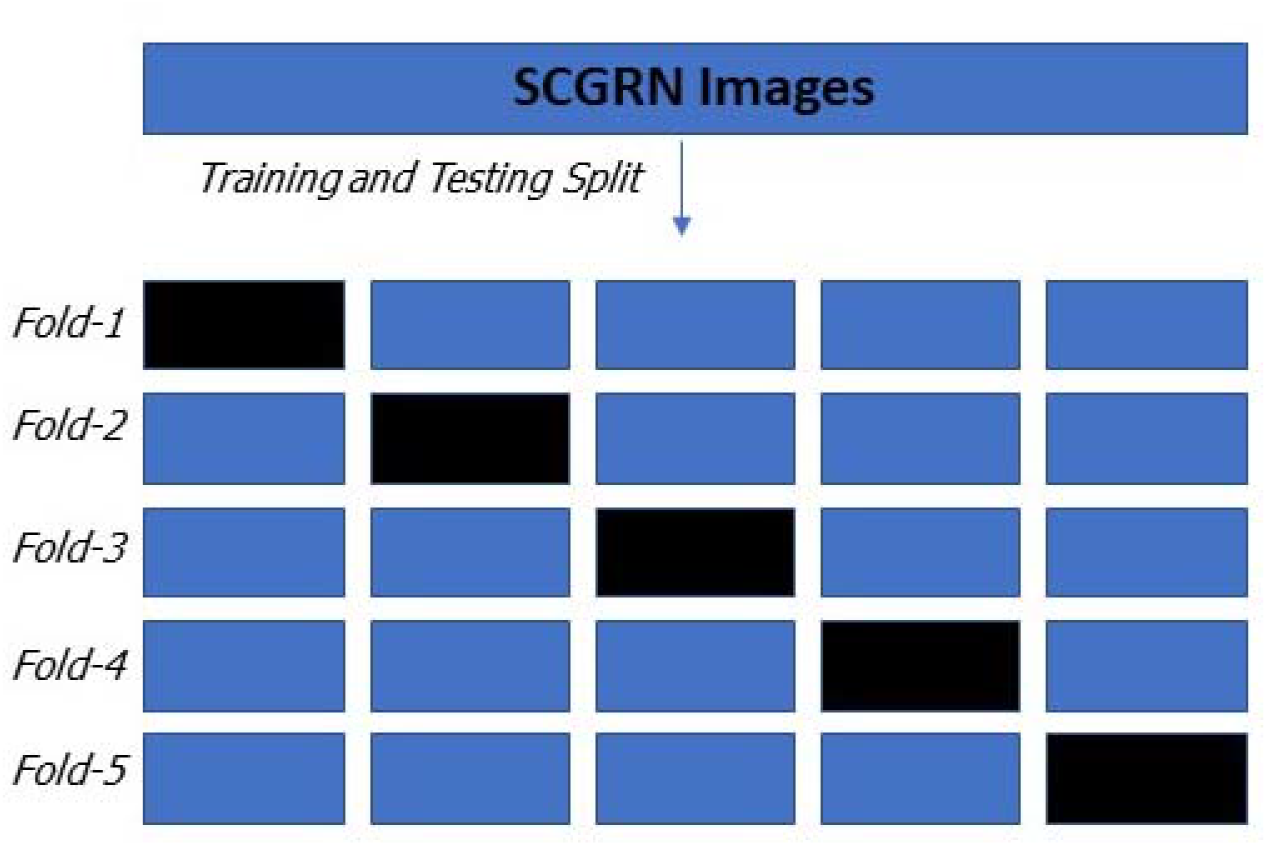
Five-fold cross-validation was performed on the whole SCGRN image dataset. Testing splits are colored in black; training splits are colored in blue.

### 3.2. Implementation Details

The DL experiments were performed using Jupyter Notebook 6.0.1, which was available with the Anaconda 4.8.1 in Python 3.7.4 [36]. To run DL architecture, we utilized Keras library. Data preprocessing and handling were performed using NumPy and Pandas libraries. Since training DL models was time-consuming and required extensive computational resources [37, 38], we ran the DL architectures on an HP workstation with a single Nvidia RTX 2080Ti GPU of 4352 CUDA cores with 11GB Memory. For evaluating all the models, we utilized Sklearn library. As per Iacono et al, we used bigSCale package in R to generate an SCGRN for healthy and T2D pancreas [19]. For reporting statistical tests using Friedman post-hoc test with Bergmann-Hommel’s procedure, we used scmamp in R [39].

### 3.3. Classification Results

Based on the 224 images of SCGRNs in healthy and T2D pancreas, obtained from human pancreatic islets, we evaluated all the DL models, thereby reporting performance on training and testing by utilizing 5-fold cross-validation.

#### 3.3.1. Training Results

Figure 5 reports the DL performance of four performance measures on the training sets during 5-fold cross-validation. Boxplots showed Xception, DenseNet121, and DenseNet169 to generate the best performance results. Specifically, they generated the highest performance of 1 or all performance measures (ACC, AUC, F1, and MCC); ResNet101 generated the lowest performance results.

**Figure 5:**
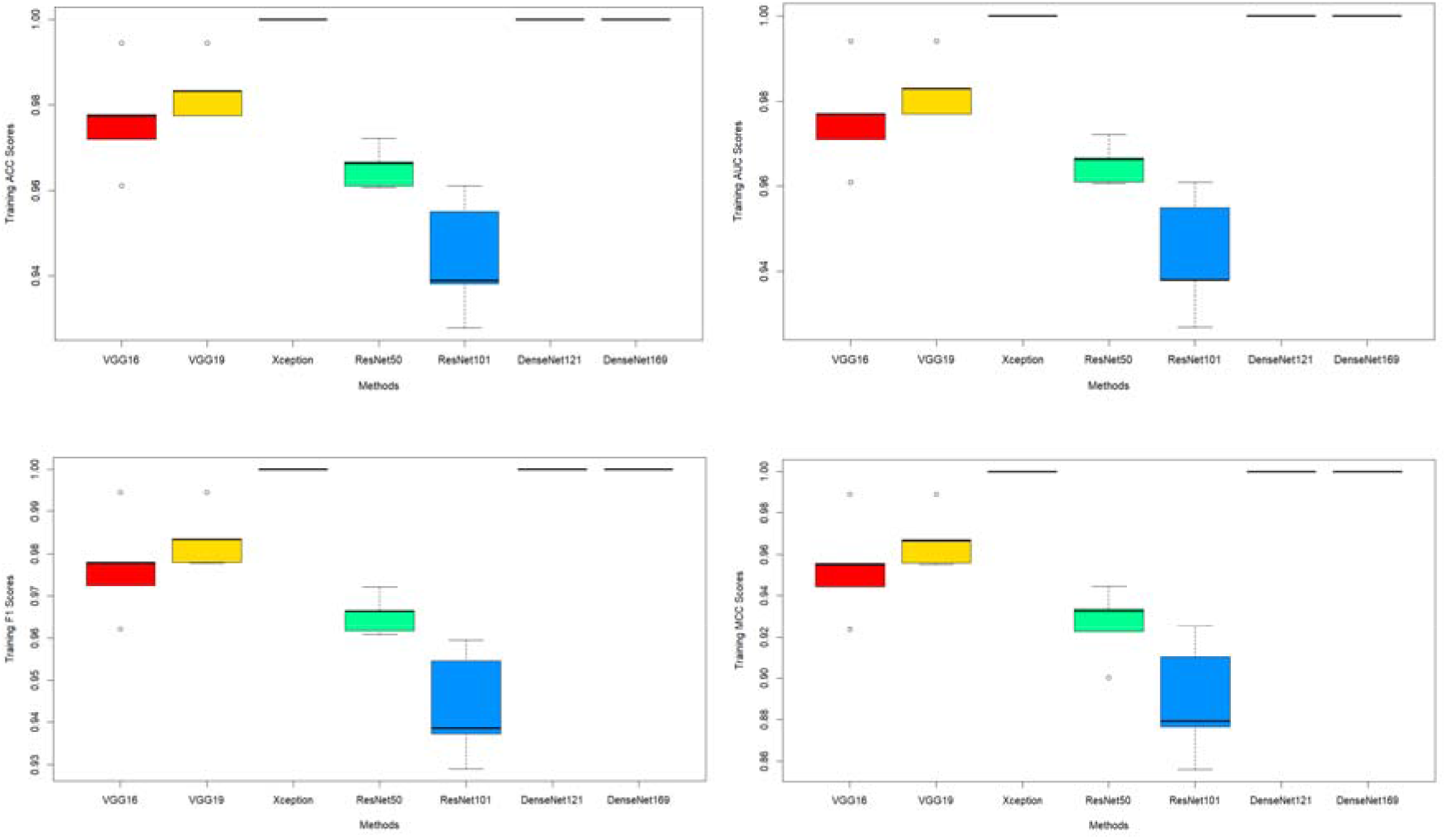
Training performance results during 5-fold cross-validation using ACC, AUC, F1, and MCC performance measures. Boxplot is used to display the performance measures of seven deep learning models.

#### 3.3.2. Testing Results

Figures 6–10 display the confusion matrices for 5 folds of test sets during crossvalidation. For each DL model, adding the numbers of all 5 confusion matrices yielded 224, which corresponded to the whole SCGRN dataset. Based on the confusion matrices in Figures 6–10, we reported in Figure 11 the performance results of DL models on the test sets during 5-fold cross-validation. Figure 11 clearly indicated VGG19 to outperform all DL models, according to ACC, AUC, F1, and MCC performance measures. For folds during cross-validation, VGG19 generated high average performance results (72–86%) based on ACC, AUC, F1, and MCC (Table 2). ResNet101 was the worst among all models, generating only average performance results (48–74%) of ACC, AUC, F1, and MCC. According to Table 2, VGG19 generated the most reliable performance results, considering all performance measures.

**Table 2:** Standard deviation (SD) of deep learning (DL) models during the 5-fold crossvalidation on test sets for ACC, AUC, F1, and MCC. MACC is Mean ACC, MAUC is Mean AUC, MF1 is Mean F1 MMCC is Mean MCC. SD stands for standard deviation. Dl with the highest mean performance measure is shown in bold.

|  | VGG16 | VGG19 | Xception | ResNet50 | ResNet101 | DenseNet121 | DenseNet169 |
| --- | --- | --- | --- | --- | --- | --- | --- |
| SD (MACC) | 0.029(0.830) | 0.041(**0.861**) | 0.046(0.777) | 0.057(0.759) | 0.029(0.740) | 0.051(0.847) | 0.050(0.839) |
| SD (MAUC) | 0.029(0.830) | 0.041(**0.861**) | 0.046(0.777) | 0.025(0.759) | 0.029(0.740) | 0.051(0.847) | 0.050(0.839) |
| SD (MF1) | 0.023(0.837) | 0.035(**0.861**) | 0.052(0.771) | 0.072(0.751) | 0.027(0.738) | 0.056(0.842) | 0.044(0.843) |
| SD (MMCC) | 0.057(0.665) | 0.081(**0.727**) | 0.089(0.560) | 0.114(0.524) | 0.058(0.487) | 0.102(0.699) | 0.101(0.682) |

**Figure 6:**
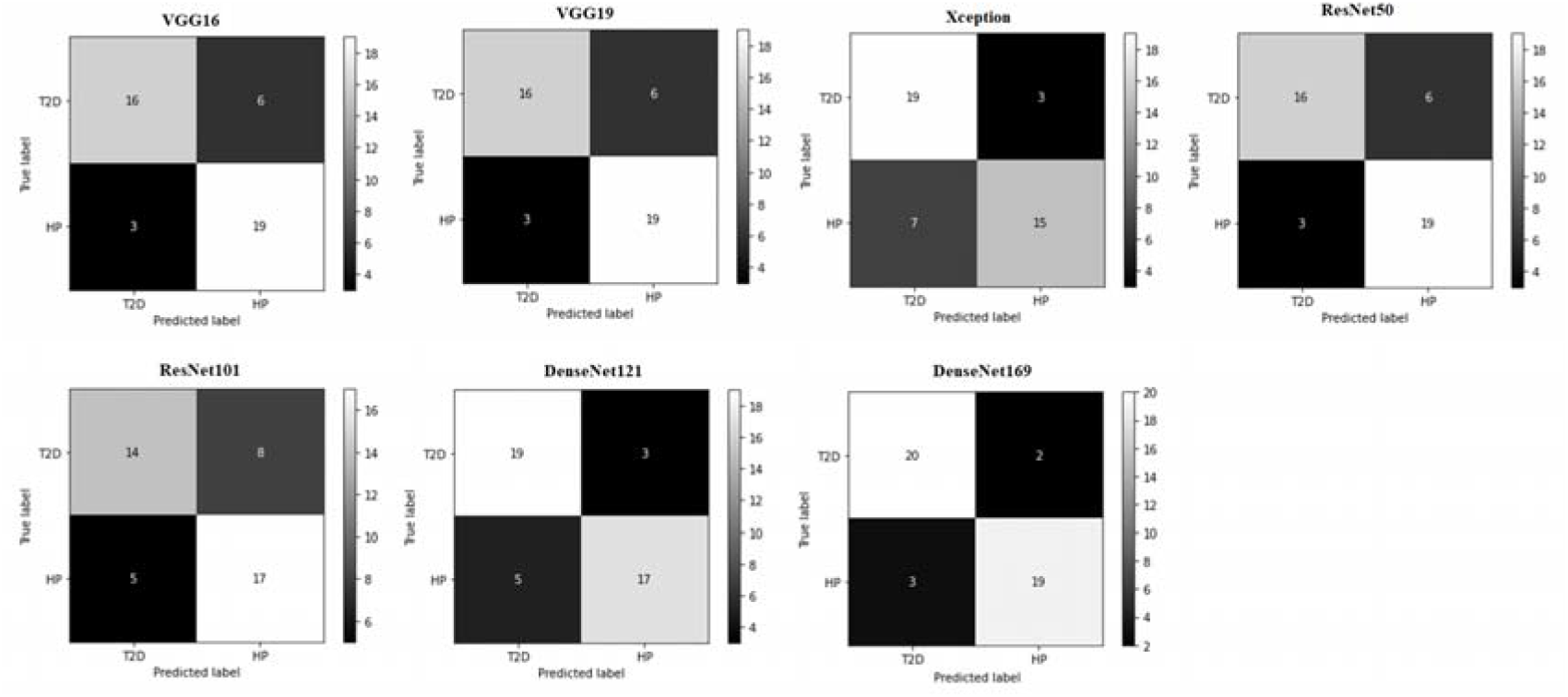
Confusion matrices for deep learning models on the test sets during Fold-1.

**Figure 7:**
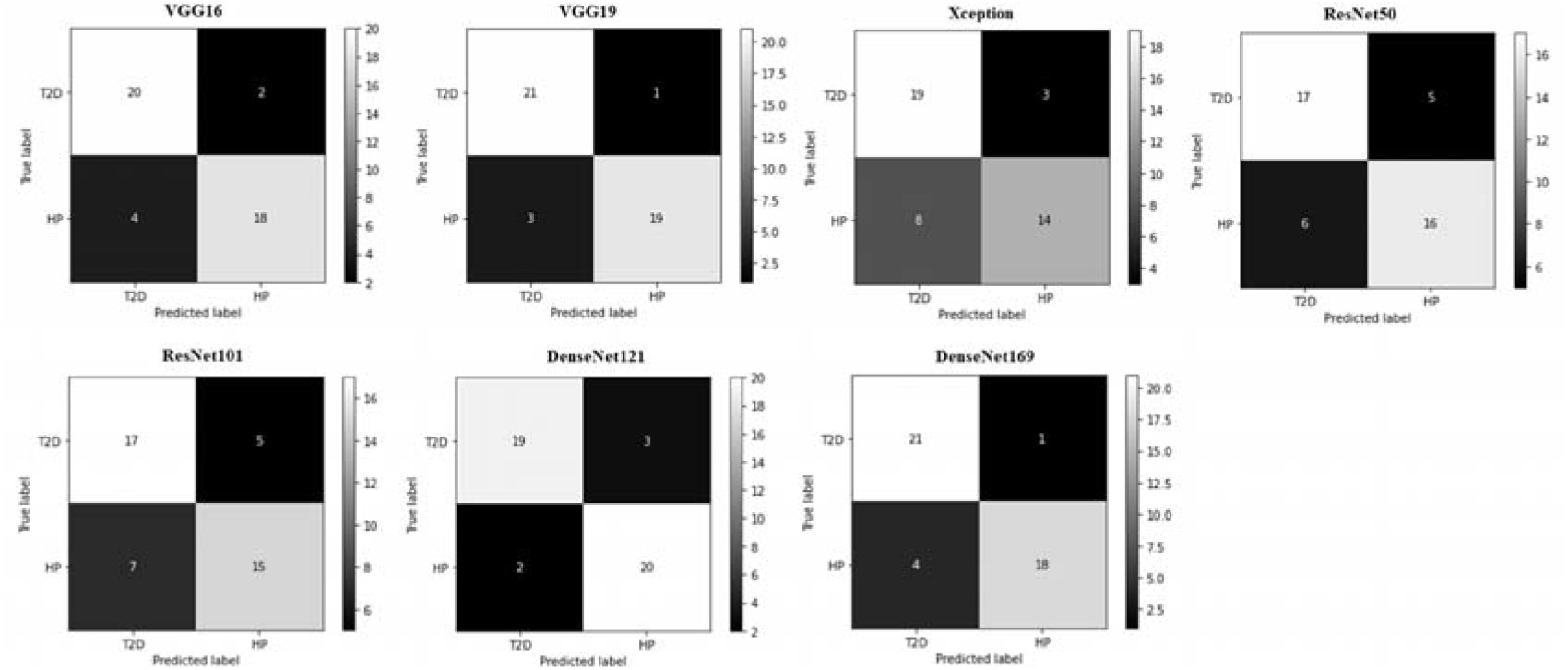
Confusion matrices for deep learning models on the test sets during Fold-2.

**Figure 8:**
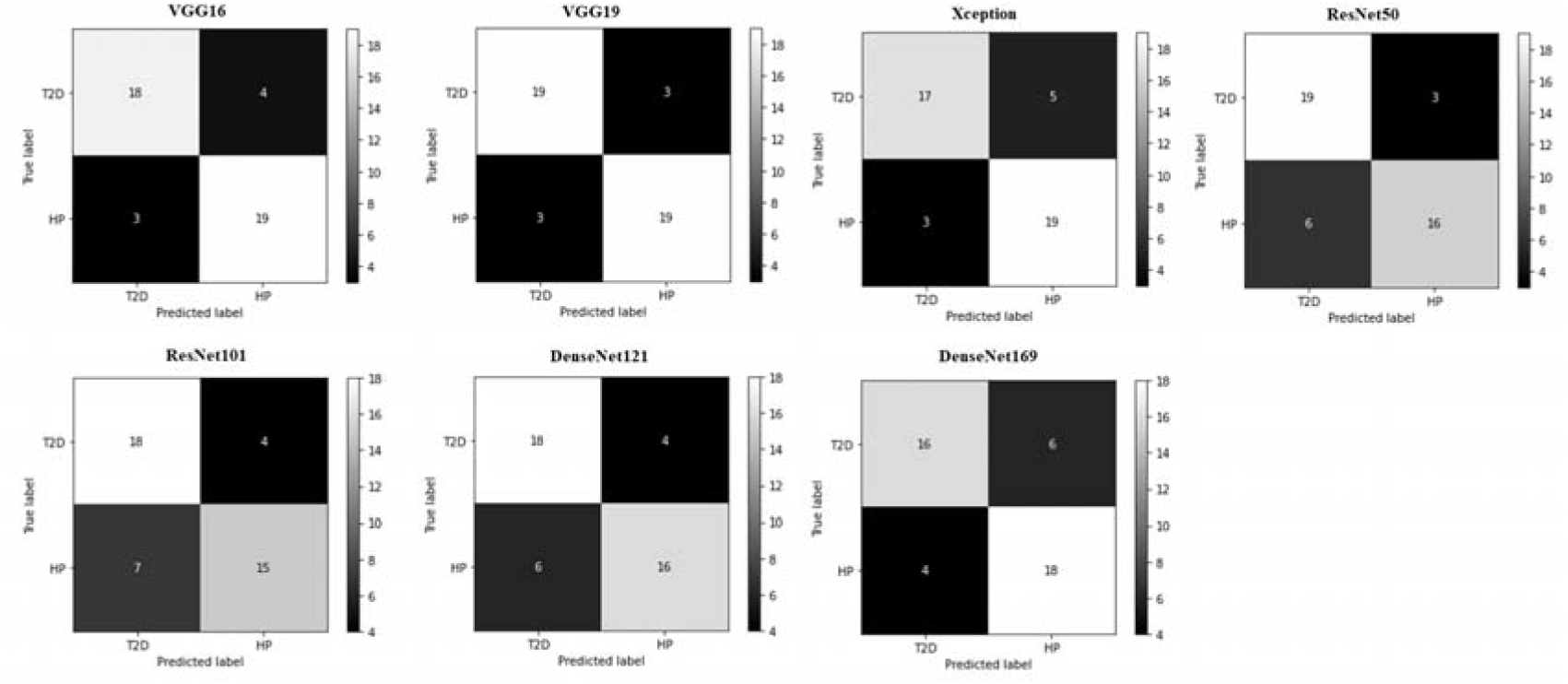
Confusion matrices for deep learning models on the test sets during Fold-3.

**Figure 9:**
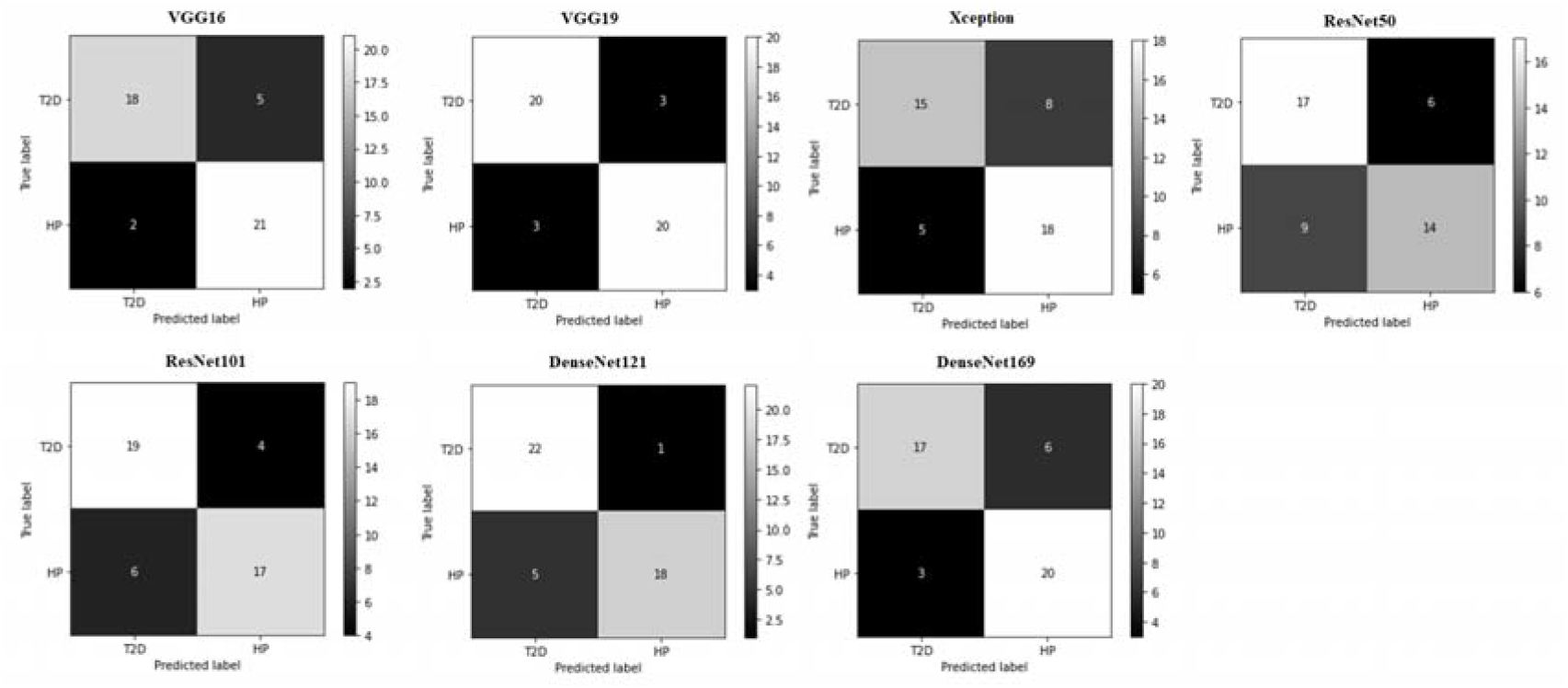
Confusion matrices for deep learning models on the test sets during Fold-4.

**Figure 10:**
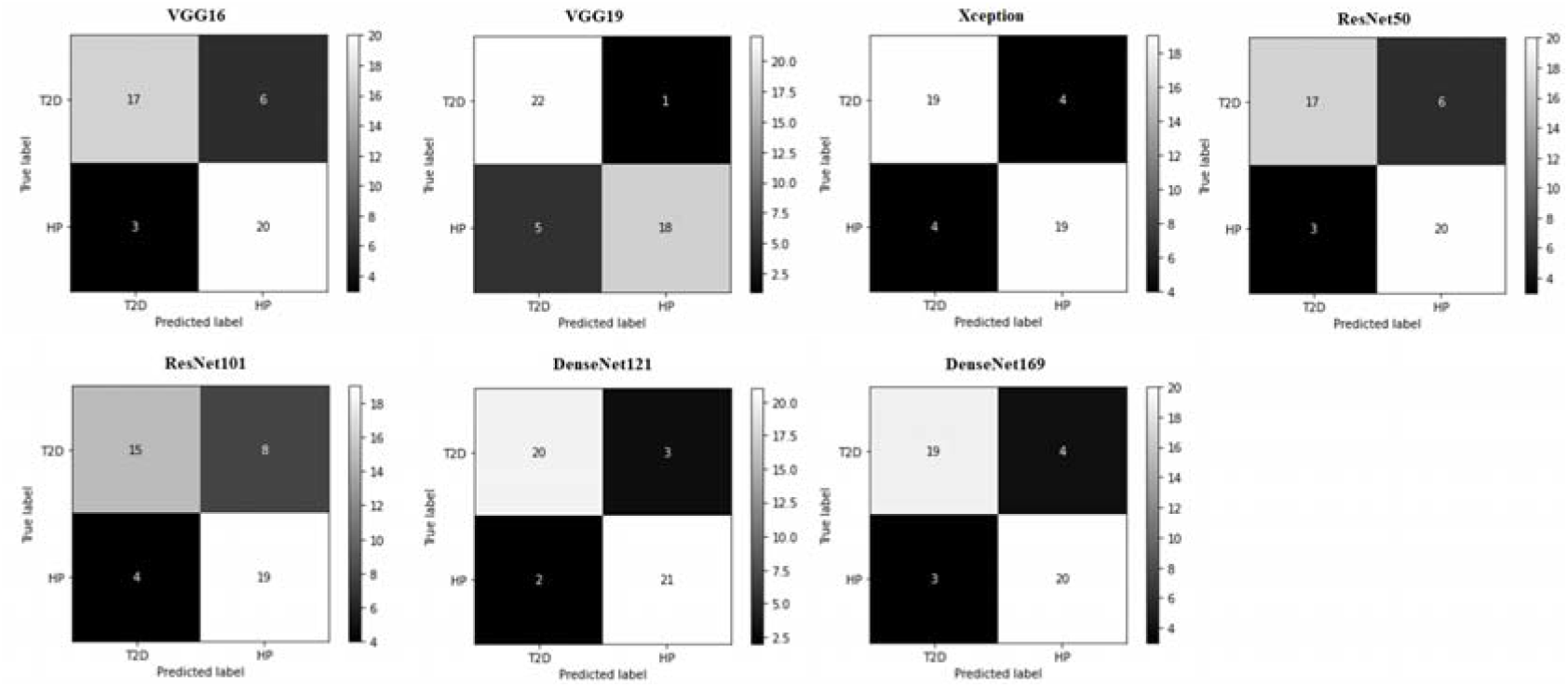
Confusion matrices for deep learning models on the test sets during Fold-5.

**Figure 11:**
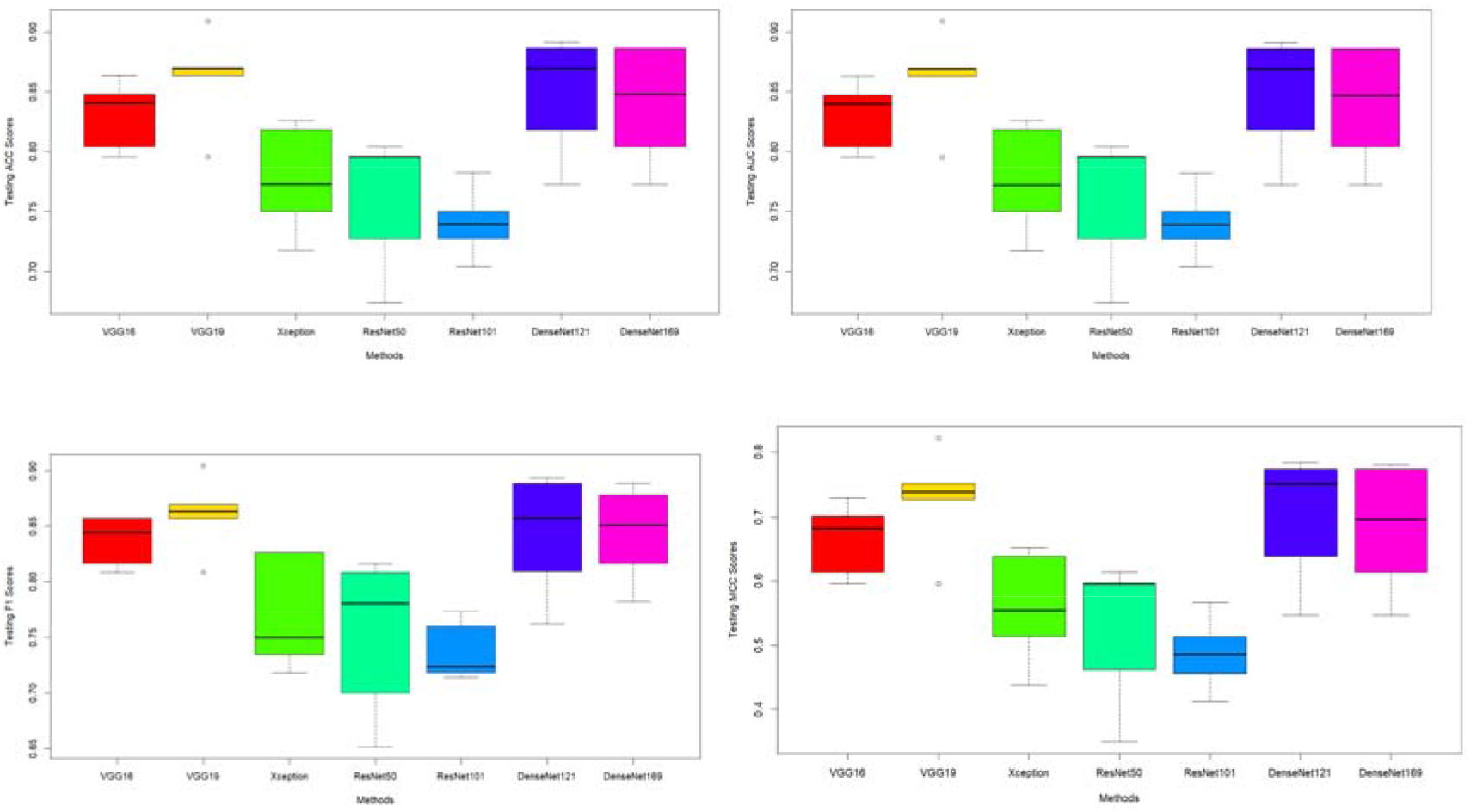
Test performance results during 5-fold cross-validation using ACC, AUC, F1, and MCC performance measures. Boxplots display the performance measures of seven deep learning models on whole datasets, consisting of 224 image data of SCGRNs.

In Figures 12a-d, we utilized Friedman post-hoc test with Bergmann and Hommel’s correction, reporting pairwise *p*-values and average performance ranking of each DL model. According to Friedman post-hoc test, applied to the results obtained from ACC and F1 (see Fig. 12a,c), VGG19 had the best average performance ranking. The Bergmann and Hommel’s corrected *p*-values showed VGG19 to significantly outperform ResNet101. However, the average performance ranking of the other DL models (i.e., VGG16, Xception, ResNet50, DenseNet121, and DenseNet169) was not significantly different from that of VGG19. The second-best average ranking was for DenseNet121. However, it was not statistically different from the other DL models. When Friedman post-hoc test was applied to results obtained from AUC and MCC performance measures, the p-values showed the average performance ranking of VGG19 and DenseNet121 to be statistically different from that of ResNet101. On the other hand, average ranking of VGG16, Xception, ResNet50, and DenseNet169 was not statistically different from that of VGG19 and DenseNet121 (see Fig. 12b,d).

**Figure 12:**
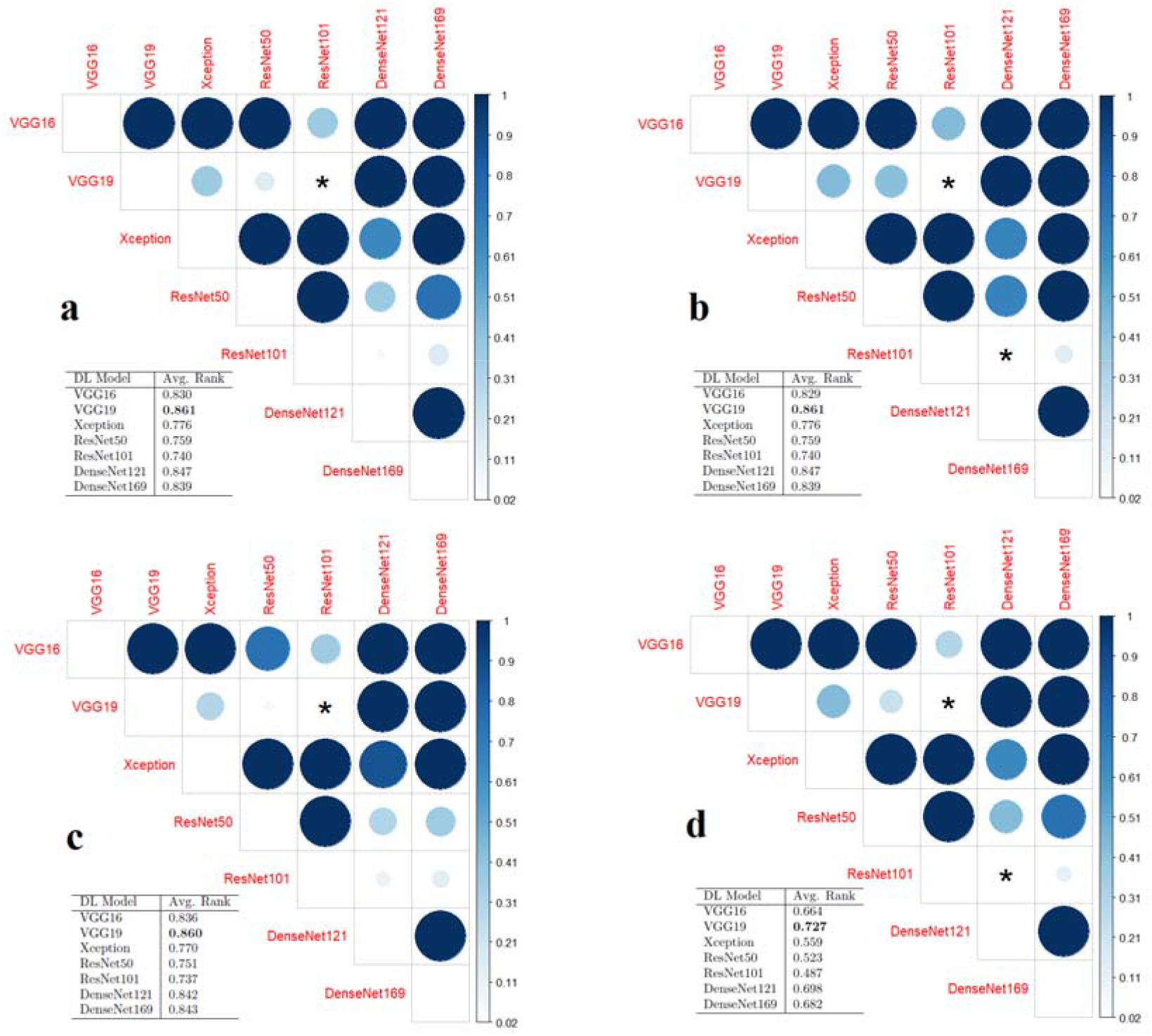
Reported *P*-values and related average rankings of all pairwise deep learning (DL) models on the test sets, based on the Friedman test with Bergmann and Hommel’s correction. Highest rank of a DL model indicated its performance to be better than that of others (shown in bold). DL models generated statistically significant results if the difference between the two models in a pair had *p* < 0.05 (shown in asterisk). Results obtained after applying the statistical test to performance results measured according to (a) ACC, (b) AUC, (c)F1, and (d) MCC.

## 4. Discussion

Our DL application consisted of three parts: (1) Generation of SCGRN image dataset from single-cell human pancreatic data, (2) formulation of the problem as a binary classification problem, where we aimed to distinguish between the SCGRNs of human pancreas with good health and with T2D, and (3) utilization and evaluation of the feasibility of several DL architectures for the image classification task. After generating our SCGRN image dataset, including 112 SCGRNs of healthy pancreas and 112 SCGRNs of T2D pancreas, we provided the image dataset to DL architecture utilizing 5-fold cross-validation. For each fold, we trained DL models on training splits and evaluated their performance on the test split.

The highly accurate predictions indicated the feasibility of using DL as an artificial intelligence tool for identifying disease differences in network biology. Utilizing such a tool, biologists would be able to (1) visualize and distinguish biological networks of complex diseases from healthy ones; and (2) improve personal experience, when facing new biological networks, by looking into similar patterns of already existing networks. In this study, several DL models generated different performance results; to show the prediction reliability across DL models during the 5-fold cross-validation, we reported standard deviation (SD) results of all models; results indicated the stability of VGG19 prediction results. Moreover, we performed a statistical test using Friedman post-hoc test with Bergmann and Hommel’s correction; the results demonstrated VGG19 to significantly outperform ResNet101 as a DL model.

To determine the factors for configuring the optimization process during the training of DL models, we use the RMSprop optimizer, setting the learning rate to 1e-5, as in a previous report [36]. The loss function used was binary_crossentropy; accuracy metric was also used. Details about loss and accuracy plots are provided in supplementary *Fold1, Fold2, Fold3, Fold4,* and *Fold5*.

In order to run computational experiments, we utilized the existing DL architectures (VGG16, VGG19, Xception, ResNet50, ResNet101, DenseNet121, and DenseNet168) for feature extraction. We provided features into a different densely connected classifier, consisting of two dense layers, as in a previous report [36]. It would be worth noting that we fine-tuned the top layers of all architectures with densely connected classifier, as described in Section 2.3. However, they did not generate good performance results. Hence, results have been placed in supplementary *Fold* files (See results within VGG16_FT,VGG19_FT, Xception_FT, ResNet50, ResNet101, DenseNet121, and DensNet169_FT).

## 5. Conclusion

We have presented a novel DL application for single-cell data analysis. First, we generated 224 images of single-cell gene regulatory networks, derived from single-cell pancreatic data, pertaining to pancreas of healthy individuals and of patients with T2D. We, then, utilized several DL architectures (VGG16, VGG19, Xception, ResNet50, ResNet101, DenseNet121, and DenseNet169) for discriminating between the SCGRNs of healthy pancreas and T2D pancreas. Each DL architecture was then trained on SCGRN images and utilized thereafter for predicting test examples. For evaluation, we utilized 5-fold cross-validation and several performance measures, including accuracy, AUC, F1, and Matthews correlation coefficient. Experimental results indicated VGG19 to generate the highest performance results. Moreover, VGG19 performed significantly better than ResNet101. We have made the 224 SCGRN image dataset freely available in supplementary *dataset*.

Future studies would be directed at: (1) Using the image dataset to devise DL models under different scenarios, such as unsupervised learning, semi-supervised learning, active learning, transfer learning, and domain adaptation, and (2) incorporating the dataset into generative adversarial networks to improve prediction performance under the transfer learning scenario [22, 40, 41].

### CRediT authorship contribution statement

TT and YT conceived and planned the research. TT conducted the deep learning experiments. TT and YT discussed the results and wrote the paper.

## Acknowledgement

This project was funded by the Deanship of Scientific Research (DSR) at King Abdulaziz University, Jeddah, under grant no. D-152-611-1440. The authors, therefore, acknowledge with thanks DSR for technical and financial support.

## Abbreviations

SCGRNs: single-cell gene regulatory networks
T2D: type 2 diabetes
DL: deep learning
DGRN: dynamic gene regulatory networks
VGG: visual geometry group
Xception: extreme inception
ResNet: residual network
DenseNet: dense convolutional network
ACC: accuracy
AUC: area under curve
MCC: Matthews correlation coefficient

**Turki Turki** received a B.S. degree in computer science from King AbdulAziz University, an M.S. degree in computer science from NYU.POLY, and a Ph.D. degree in computer science from the New Jersey Institute of Technology. He is currently an assistant professor with the Department of Computer Science, King Abdulaziz University, Saudi Arabia. His research interests include Artificial Intelligence (Tensor Learning, Machine Learning, Deep Learning) and Bioinformatics. His research studies have been published in journals such as Expert Systems with Applications, Frontiers in Genetics, Current Pharmaceutical Design, Computers in Biology and Medicine, and Genes. Dr. Turki has served on the program committees of several international conferences and is currently a review editor for *Frontiers in Artificial Intelligence* and *Frontiers in Big Data*. In addition, he is an editorial board member of *Computers in Biology and Medicine* and *Sustainable Computing: Informatics and Systems*.

**Y-h. Taguchi** received a B.S. degree in physics from the Tokyo Institute of Technology and a Ph.D. degree in physics from the Tokyo Institute of Technology. He is currently a full professor with the Department of Physics, Chuo University, Japan. His works have been published in leading journals such as *Physical Review Letters, Bioinformatics*, and *Scientific Reports*. His research interests include bioinformatics, machine learning, and nonlinear physics. He is also an editorial board member of *Frontiers in Genetics, PloS ONE, BMC Medical Genomics, Medicine* (Lippincott Williams & Wilkins journal), *BMC Research Notes*, and *IPSJ Transaction on Bioinformatics*.

